# Injectable therapeutic organoids using sacrificial hydrogels

**DOI:** 10.1101/2020.01.27.922112

**Authors:** Ninna S. Rossen, Priya N. Anandakumaran, Rafael zur Nieden, Kahmun Lo, Wenjie Luo, Christian Park, Chuqiao Huyan, Qinyuouen Fu, Ziwei Song, Rajinder P. Singh-Moon, Janice Chung, Jennifer Goldenberg, Nirali Sampat, Tetsuhiro Harimoto, Danielle Bajakian, Brian M. Gillette, Samuel K. Sia

## Abstract

Organoids, by promoting self-organization of cells into native-like structures, are becoming widespread in drug-screening technologies, but have so far been used sparingly for cell therapy as current approaches for producing self-organized cell clusters lack scalability or reproducibility in size and cellular organization. We introduce a method of using hydrogels as sacrificial scaffolds, which allow cells to form self-organized clusters followed by gentle release, resulting in highly reproducible multicellular structures on a large scale. We demonstrated this strategy for endothelial cells and mesenchymal stem cells to self-organize into blood-vessel units, which were injected into mice using hypodermic needles, and observed in real time to rapidly form perfusing vasculature. As cell therapy transforms into a new class of therapeutic modality, this simple method – by making use of the dynamic nature of hydrogels – could offer high yields of self-organized multicellular aggregates with reproducible sizes and cellular architectures.

## Introduction

Organoids, such as vascularized organoids or spheroids ^1–3^, are three-dimensional multicellular clusters which mimic the structure and function of native tissues and are useful for on-chip drug screening ^4,5^. For use as a cell therapy, delivery of cells within well-controlled microenvironments, rather than suspensions of isolated cells, could promote and maintain desired cellular functions within dynamic and complex *in vivo* environments ^6–11^. As organoids are increasingly being explored for *in vivo* studies and therapy, there is increasing recognition of the unmet challenge in generating multicellular aggregates with high reproducibility and control. As one example, even though control over “organoid size, shape, cellular composition and 3D architecture…is essential in order to understand the mechanisms that underlie organoid development in normal and pathological situations, and to use them as targets for manipulation or drug testing”, reproducibility has been cited as “the major bottleneck of current organoid systems” ^12^.

The major current methods for generating organoids include spinner cultures ^13^, hanging drops ^14,15^, and non-adhesive 96-well plates ^16–19^ (Supplementary Table 1), but these methods are difficult to scale or harsh to cells. (Alternatively, microtissues that are “cells in gels” ^20–23^ typically feature cells moving to pre-formed pores within a hydrogel scaffold, but the cells are limited in their ability to self-organize into desired structures ^24^, and the resultant gels exhibit variable structures and sizes dependent on the pores and may be undesired in the implanted site due to potential immunogenicity). More recently, methods to fabricate organoids based on micro-sized wells have faced challenges of either high adsorption (of steroid hormones, small molecules, and drugs ^25,26^ for PDMS-based wells) or inefficient and harsh processes, usually involving vigorous pipetting or high-speed centrifugation, to separate and remove the cellular clusters from the microwells. Such procedures produce cellular clusters at a low yield and could damage cellular structures and function. Recognizing this limitation, other studies have proposed more complex methods to actively release cellular clusters from microwells ^27–29^. As such, there still lacks reliable methods to generate organoids at high yield and with reproducibility and control over aggregate size and cellular organization.

Many hydrogels are biocompatible and have been used as a dynamically responsive biomaterial (such as microfluidic valve ^30^, changing cellular microenvironment ^31^, and stimuli-responsive drug release ^32^). We hypothesize that a dynamic change in the cross-linking state of hydrogels could gently release organoids, and sought to demonstrate the strategy for producing large numbers of vascularized organoids with high reproducibility and scalability, as well as the ability to retain functionality after passing through needles to obviate invasive surgery ^33,34^. We also assessed the ability of the pre-formed blood-vessel units, after injection, to rapidly integrate with the host’s vascular network in a healthy mouse model.

## Results

### Hydrogels as a sacrificial scaffold as a gentle and scalable method for producing and harvesting organoids

Sacrificial materials are widely used in micromachining of microelectromechanical systems (MEMS) to release patterned metals or semiconductors from a substrate (Figure 1a). To cellular structures, some hydrogels (such as agarose or poly(ethylene)glycol as previously demonstrated) can be non-adhesive and thereby promote cells to interact with one another and contract into microtissues and organoids ^1,35^. We hypothesized that a dynamic change in the cross-linking state of alginate, which can be achieved by adding calcium or a chelator and has been demonstrated for other purposes ^31,36^. could similarly release cell-based structures from a surface without significantly disrupting the organoid structures or underlying cell function ^37,38^. Specifically, we deposit the sacrificial material, create the sacrificial structure by cross-linking the alginate in its patterned state, deposit cells on top to allow cellular self-organization to take place, and remove the sacrificial layer by adding a chelator (5% w/v sodium citrate) (Fig. 1a; see Supplementary Figure 1 for fabrication details). The alginate is uncrosslinked within ∼12 minutes (Fig. 1b), to gently release a large number of organoids floating in solution (Fig. 1c,d; Supplementary Video 1). The resulting organoid solution could be gently pipetted into tubes, centrifuged and resuspended in a culture medium suitable for downstream manipulation or direct cell delivery. Each step is simple, can be conducted with sterile liquid handling, and can be automated.

**Fig. 1.**
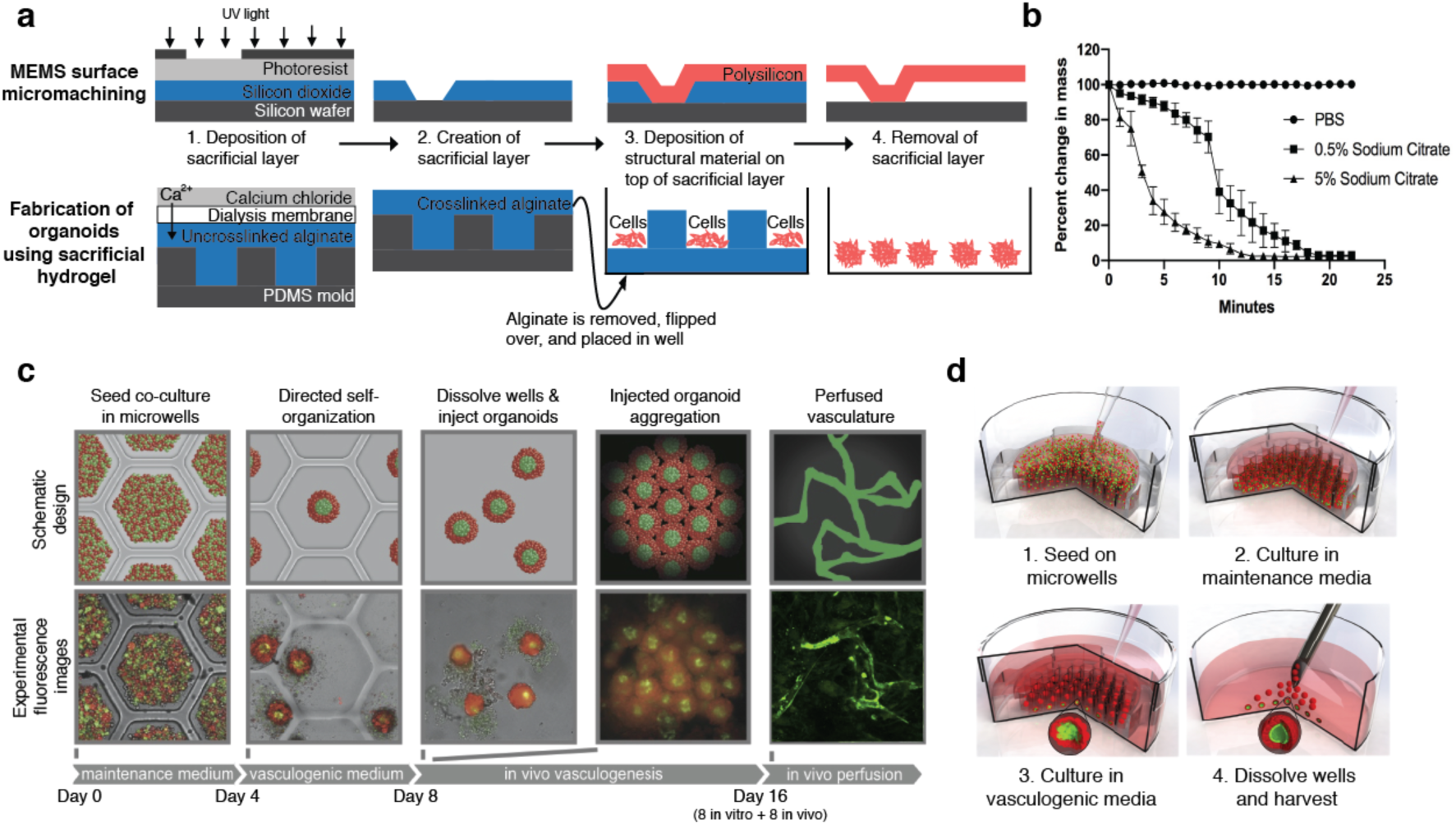
Schematic diagram of method of using sacrificial hydrogels to produce therapeutic organoids. **(A)** Schematic demonstrating the parallels between the surface micromachining method to fabricate MEMS devices such as a microcantilever (top) and the use of sacrificial alginate microwells to fabricate organoids (bottom). Both methods involve the use of a sacrificial layer (blue) to fabricate the final structure (red). **(B)** Time required to completely uncrosslink alginate microwells following incubation with different concentrations of a chelator (sodium citrate) by measuring the percent change in mass over time (n=3, Error bars are standard deviations). **(C)** Schematic diagrams (top) and corresponding experimental images (bottom) showing the steps of organoid fabrication and *in vivo* perfusion. Experimental data were collected using GFP-labelled HUVECs and RFP-labelled mouse MSCs. First, a co-culture of endothelial cells (green) and therapeutic cells (red) is seeded on dissolvable alginate microwells. Second, after being cultured in maintenance medium without growth factors for 3 to 4 days, cells self-organize into organoids with an endothelial core. A switch into culture medium with vasculogenic growth factors for an additional 4 days promoted formation of vessels within the organoids. Third, alginate microwells were dissolved with 5% sodium citrate to release organoids. Fourth, suspension of organoids could be centrifuged and assembled into a macro-tissue *in vitro* to study vascular formation, or injected into the subdermis or ischemic hindlimb of a mouse to demonstrate engraftment *in vivo*. Fifth, injected organoids rapidly connected to form perfused microvasculature *in vivo*. **(D)** The liquid handling steps in the process: 1) seeding the co-culture of ECs (green) and MSCs (red) by pipetting cells onto alginate microwell construct, 2) adding maintenance media once the cells have settled to the bottom of the microwells (approx. 30 minutes), 3) switching to vasculogenic media once an endothelial core has formed, and 4) gently dissolving the alginate microwells (approx. 5 minutes) to harvest organoids (the organoids can be gently washed prior to injection).

The step of cellular self-organization can be adjusted depending on the organoid of interest. For vascularized organoids (Fig. 1c), we seeded a co-culture containing ECs and MSCs (of either mouse or human origin) into the dissolvable alginate microwells. We cultured the cells in media without growth factors (“maintenance” medium), followed by a vasculogenic medium with growth factors to induce cell-cell interactions including sprouting of blood vessel-like structures. The organoids, now containing blood vessel-like structures, contract and are gently released by dissolving the alginate microwells.

The same small number of steps (Fig. 1c, d) can harvest a large number of organoids by using alginate templates with large numbers of microwells. As an example, we demonstrated three different sizes of alginate microwell inserts for culture dishes (Fig. 2a): 15.6 mm-diameter inserts containing >1000 microwells (yielding >24,000 organoids on a 24-well plate), 22.1-mm inserts containing >3000 microwells (yielding >36,000 organoids on a 12-well plate), and a 60-mm diameter insert containing >30,000 microwells in a 60-mm culture dish. If desired, the inserts can be stacked to increase the number of organoids produced in the same area with additional media changes.

**Fig. 2.**
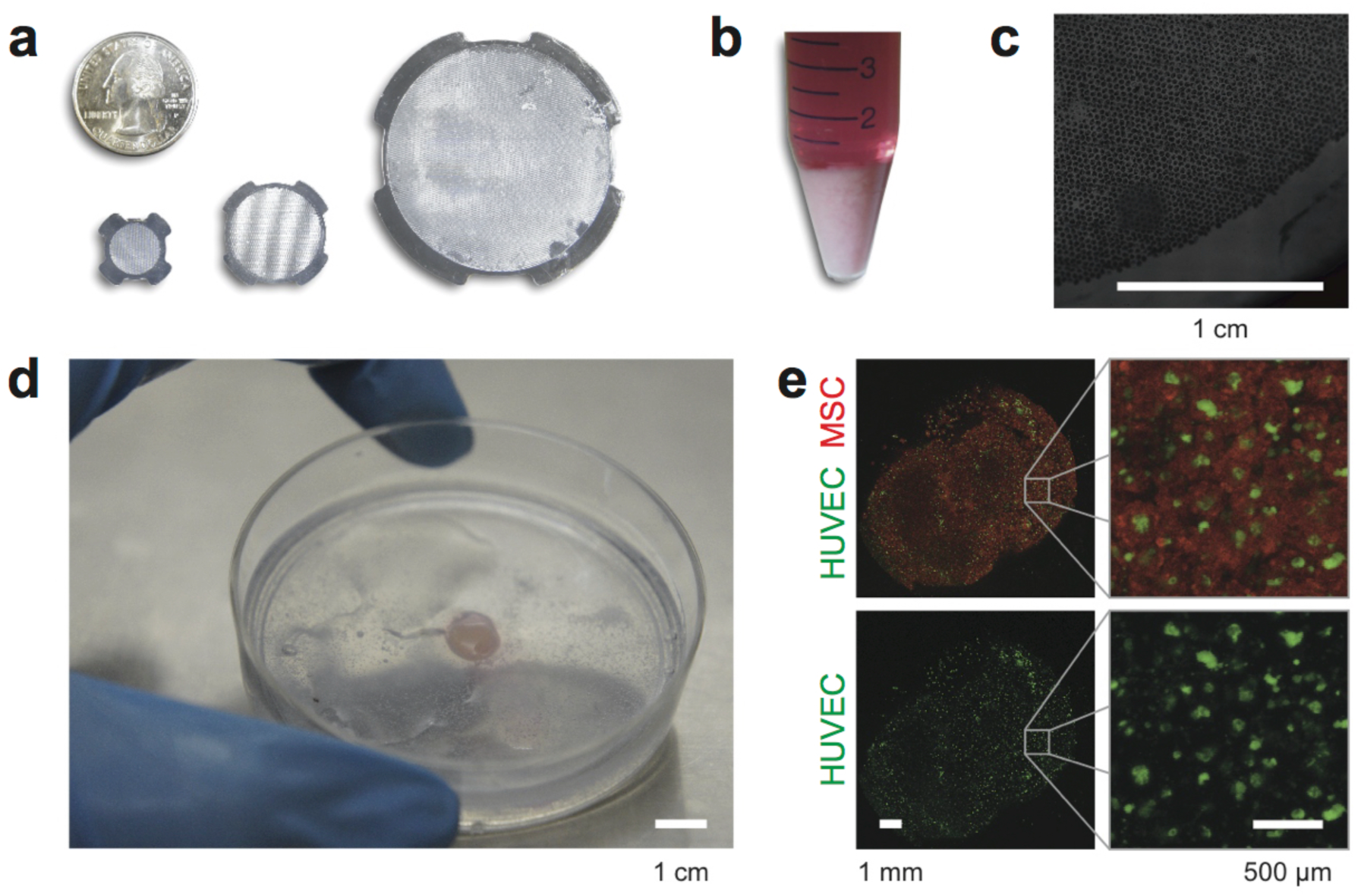
Production of organoids at large scale, and functionality of organoids to form macrotissue. **(A)** Pictures of three alginate microwells constructs for inserts into 24-well plates, 12-well plates or 60-mm dishes with the capacity to produce 24 × 1000, 12 × 3000 or 30,000 organoids respectively. **(B)** Picture of 250 million cells for seeding into alginate microwells. Cells in this figure are GFP-labelled HUVECs and RFP-labelled mouse MSCs. **(C)** Stitched brightfield image of cells seeded in a 60-mm construct with 30,000 wells to create 30,000 organoids. Scale bar is 1 cm. **(D)** Picture of a 1 mm thick macrotissues with an area of 1 cm^2^ assembled *in vitro* by collecting the 30,000 mature pre-vascularized organoids produced with the alginate microwell (a and b) construct in a 60-mm dish. Scale bar is 1 cm. **(E)** Fluorescence images of the macrotissue in (d) with a close-up of the closely packed organoids with endothelial cores (green). Scale bars are 1 mm (left) and 500 µm (right).

We demonstrated this massive parallel production of more than 30,000 organoids by seeding a quarter of a billion cells (Fig. 2b) in one 60-mm dish insert (Fig. 2c). We also tested the ability of the organoids to assemble *in vitro* into a microvascular network (Fig. 2d). We co-cultured RFP-labelled MSCs and GFP-labelled ECs for four days, gently harvested the organoids, and assembled them into a macroscopic tissue with surface area of 1 cm^2^ and a height of 1 mm (Fig. 2d). We performed fluorescence imaging of this macroscopic tissue (Fig. 2e). The organoids were densely packed, and exhibited distinct endothelial core structures, confirming that the gentle harvest and assembly did not disturb the internal architecture of the organoids. The assembled macrotissue, consisting of fully contracted organoids, did not visibly contract during subsequent *in vitro* culture.

### Production of organoids with reproducible size and structure

Next, we studied whether the sizes and internal architectures of organoids could be controlled reproducibly. In the absence of exogenous growth factors, we observed that GFP-labelled human umbilical vein endothelial cells (HUVECs), which were initially randomly distributed alongside RFP-labelled mouse MSCs, migrate to the center of the organoids and form endothelial cores after culture in the “maintenance” medium for 3 days (Fig. 3a, top panel, and Supplementary Video 2). Similarly, ECs also formed endothelial cores when co-cultured with another cell type (fibroblasts) in a medium without growth factors (Supplementary Figure 2), the endothelial cores were more pronounced than in a previous observation ^19^. By contrast, ECs did not migrate to the center when the organoids were initially cultured in a vasculogenic medium containing 50 ng/mL VEGF and 50 ng/mL bFGF (Fig. 3a, bottom panel), consistent with a previous observation ^9^. Overall, the data showed the organoids to exhibit reproducible internal architectures containing endothelial cores.

**Fig. 3.**
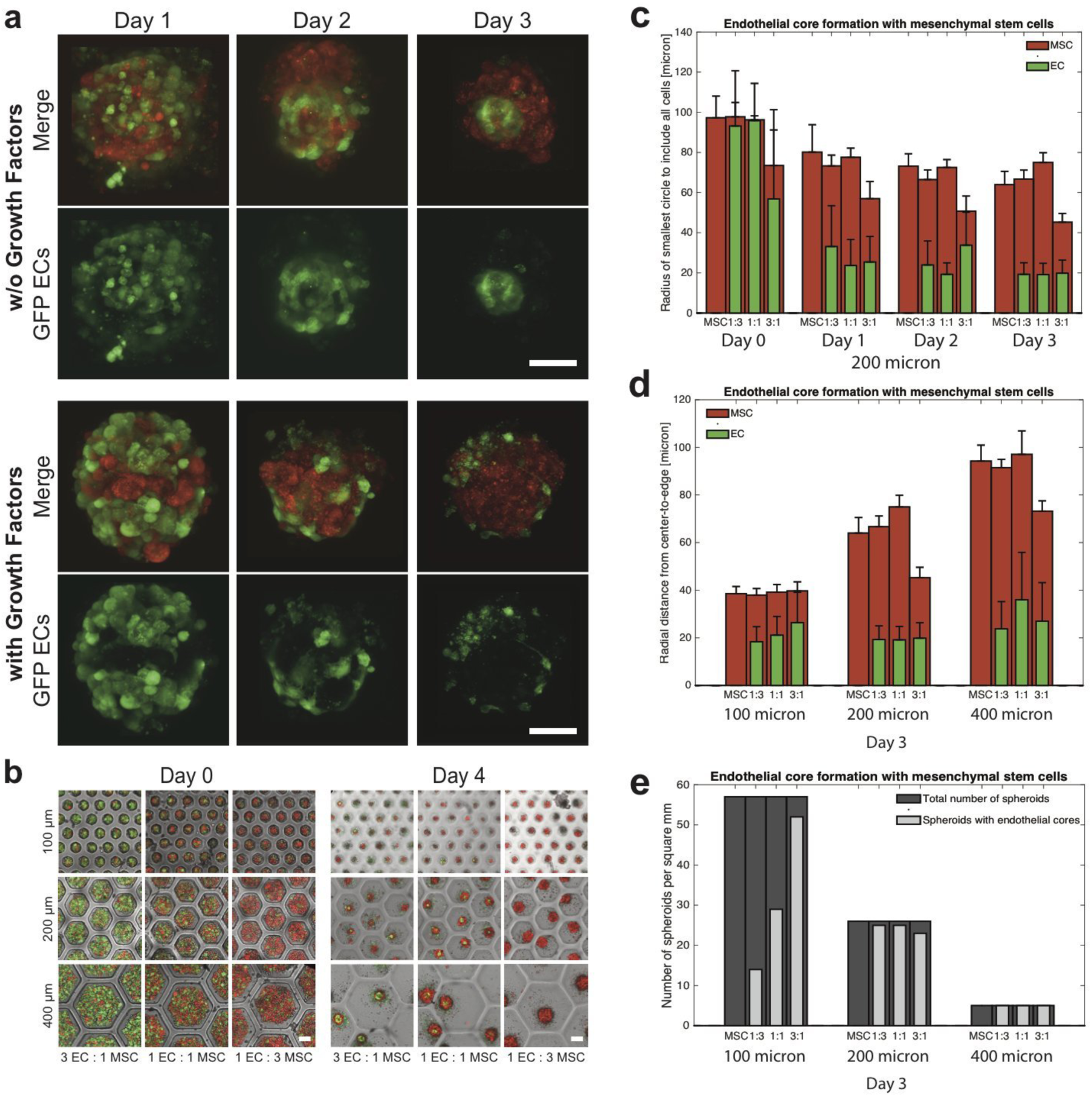
Production of vascularized organoids with high reproducibility in size and structure. **(A)** Confocal fluorescence images of co-culture organoids of GFP-labeled HUVECs (green) and RFP-labeled mouse MSCs (red) over the first three days in maintenance medium without growth factors (top) or in vasculogenic medium with 40 ng/mL VEGF and 40 ng/mL bFGF (bottom). The cells self-organize by migration, and either formed endothelial cores when cultured in media without growth factors (top) or had endothelial cells randomly distributed near the surface of the organoid and did not form endothelial cores when cultured in media with growth factors (bottom). Scale bars are 100 µm. **(B)** Overlay of fluorescent and transmitted images showing parallel production of organoids in arrays of different sizes of microwells (with either 100, 200, or 400 µm diameter) and different co-culture ratios (1 EC : 3 MSC, 1 EC : 1 MSC or 3 EC : 1 MSC). Different sizes of microwells yield different sizes of organoids, either unvascularized with only MSCs or pre-vascularized with a co-culture of ECs and MSCs, and different co-culture ratios yield different endothelial core sizes. Scale bars are 100 µm. **(C)** Quantitative analysis of cell aggregation into organoids and the formation of an endothelial core over time in 200 µm microwells, as measured by the radius of the smallest circle that can contain all MSCs (red) or all ECs (green) (*n* > 20). **(D)** Barplot showing the size of fully-contracted organoids (red) and the size of the endothelial cores (green) for all tested microwell sizes and co-culture ratios. **(E)** Reproducibility of endothelial cores; the number of organoids produced in 1 mm^2^ (dark grey) and the number of organoids containing and endothelial core (light grey) for all tested microwell sizes and co-culture ratios.

We characterized the reproducibility of the method in controlling the size of the pre-vascularized organoid. By varying the microwell sizes and the co-culture ratios of cell types (Fig. 3b), we controlled the number of cells that could aggregate into a single organoid. For example, microwells of three different sizes (100, 200, and 400 µm diameter) yielded organoids of three different sizes (39±3 µm, 71±5 µm, and 82±7 µm diameter, respectively, all at the same cell-seeding concentration) (Fig. 3b). The well size was chosen to be large enough to hold all the cells at the initial seeding concentration, but small enough to ensure sufficient cell-cell contact to form a single organoid rather than multiple organoids. The cells aggregated into compact organoids within the first two days of *in vitro* culture, as seen by the decreasing radius of the smallest circle to include all cells (Fig. 3c), with the main contraction happening in the first day and no further contraction after three days. We also observed that the size of the fully contracted organoids (at day 2 and after) correlated to the number of cells in the organoid as expected; the diameter of the organoids’ cross sections related to the number of cells in the organoid and the cells typical volume as r_organoid_ = (6/π v_cell_ n_cell_)^1/3^/2 (Supplementary Figure 3).

Also, we quantitatively analyzed the formation of organoids for cultures containing only MSCs, and co-cultures with EC:MSC ratios of 1:3, 1:1, and 3:1 (Fig. 3b-e). In 200-µm microwells, over three days, cells contracted into an organoid and ECs migrated towards the center (Fig. 3c), and co-cultures in 400 µm microwells showed similar trends in organoid contraction and EC migration (Supplementary Figure 4). Co-cultures in 100-µm microwells, however, did not contain enough cells (fewer than 150 cells in total) to form a distinct center (Supplementary Figure 5). We also observed that the organoids per unit area and the number of organoids containing defined internal architectures could be controlled by varying microwell sizes and ratios of cell types (Fig. 3e). (In subsequent *in vivo* studies, we have used 200-µm microwells with ratios of MSC only, 1 EC:3 MSC and 1 EC:1 MSC, as these conditions showed aggregation involving almost all the cells within the microwells.) Overall, the data showed the method can produce organoids with internal architectures at high throughput and different sizes controllably.

### Production of pre-vascularized human organoids with reproducible size and structure

We examined the effectiveness of this method for producing pre-vascularized organoids containing human adipose-derived MSCs (hAMSCs) with human umbilical vein endothelial cells (HUVECs), in ratios of MSCs only, 1 EC:3 MSC, and 1 EC:1 MSC. We examined the maturation of organoids over 8 days, where organoids were first grown in maintenance medium over 3 days to form endothelial cores, and then switched to vasculogenic medium containing exogenous growth factors for 5 days (Fig. 4a). By day 8, vessel-like structures, such as lumens within the center of the organoid, with sprouting and maturation of vessels towards the surface were observed (especially evident in the larger organoids of the 400-µm wells). The initial migration of ECs was apparent after 20 hours (Fig. 4b and Supplementary Figure 6). In addition, we placed multiple pre-vascularized organoids inside 400 µm alginate wells that were collagen-doped, to mimic the adhesiveness of native tissues. Within 24 hours, organoids attached to each other and contracted to form a larger, compact mesotissue (aggregation of multiple organoids) with a smooth outer border (Fig. 4c, with additional time points in Supplementary Figure 7 and Supplementary Video 3). Hence, this method produced organoids containing human ECs and MSCs, with control over sizes and spatial architectures, and confirming the ability to form a pre-vascularized mesotissue.

**Fig. 4.**
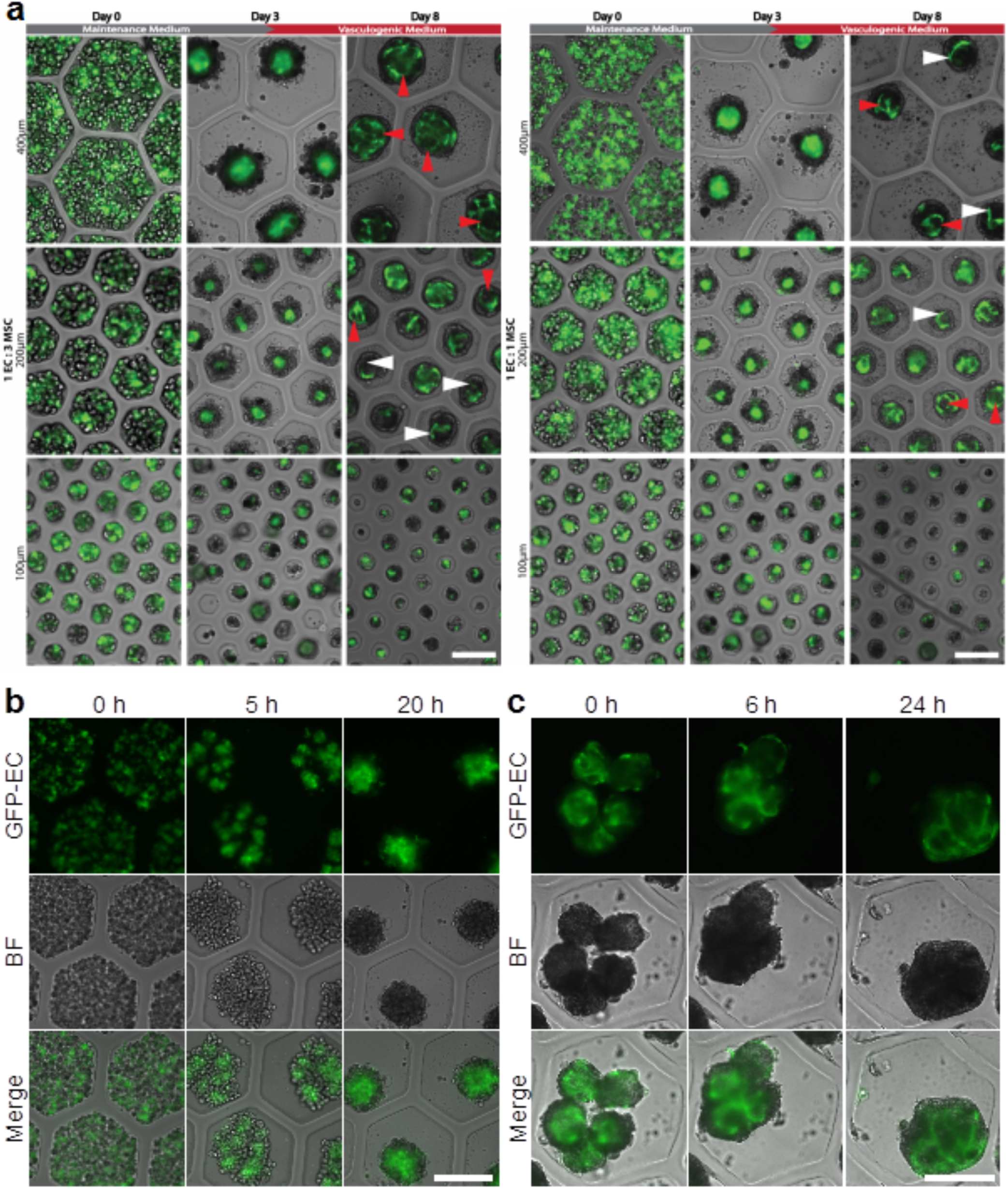
Production of vascularized organoids with human cells. **(A)** Maturation of endothelial cores with dynamic culture conditions for two co-culture ratios; 1 GFP-HUVEC : 3 hAMSC (left) and 1 GFP-HUVEC : 1 hAMSC (right). The cells are seeded (day 0) and initially cultured in maintenance medium without growth factors to form endothelial cores. After 3 days the organoids were cultured in vasculogenic medium with 40 ng/mL VEGF and 40 ng/mL bFGF and the endothelial cores matured into vessels with discernable lumens (red arrows) and sprouts (white arrows). Scale bars are 200 µm. **(B)** Epifluorescence, brightfield, and overlay images showing early self-organization of pre-vascularized organoids over the first 20 hours, with a 1 GFP-HUVEC : 1 hAMSC co-culture in 400 µm microwells. Scale bar is 200 µm. **(C)** Epifluorescence, brightfield, and overlay images showing fusion of pre-vascularized organoids (same conditions as in right a and b) into mesotissues over the first 24 hours of the fusion process within a 400 µm collagen-doped alginate microwell. Scale bar is 200 µm.

### Rapid host perfusion of pre-vascularized organoids in mouse model

Next, we assessed the effectiveness of the prevascularized organoids to self-organize to form a vascular network, anastomose to native host vasculature, and be perfused with host blood in a mouse model (Fig. 5a). To facilitate real-time visualization, we performed surgery to place a window chamber (Supplementary Figure 8) to permit brightfield, epifluorescence and confocal imaging. We used organoids formed in 200-μm wells yielding organoids approximately 70 μm in diameter, which is also within the diffusion limit of oxygen ^39^. We produced and harvested pre-vascularized organoids made of human cells (HUVECs and hAMSCs), which we injected into SCID mice, a well-established animal model for studying integration of xenografts made of human cells ^40^. We could inject and monitor the vascular formation for multiple different conditions (e.g. 1 HUVEC : 1 hAMSC and hAMSC only) in the same mouse, by utilizing the strong bond between the fascia and the subdermis. We injected the organoids through the fascia and into the space between the fascia and the subdermis, leaving the subcutaneous tissue intact between injection sites to create a barrier (Fig. 5b). The organoids held up intact to the shear stress of injection through a syringe and needle (Supplementary Figure 9). Interestingly, the shell of MSCs shielded the central blood-vessel building block against shear, and preserved the organoids’ architectural integrity after they passed through the needle. (We also demonstrated the organoids could be injected directly into adipose tissue (Supplementary Figure 10) with good integration.)

**Fig. 5.**
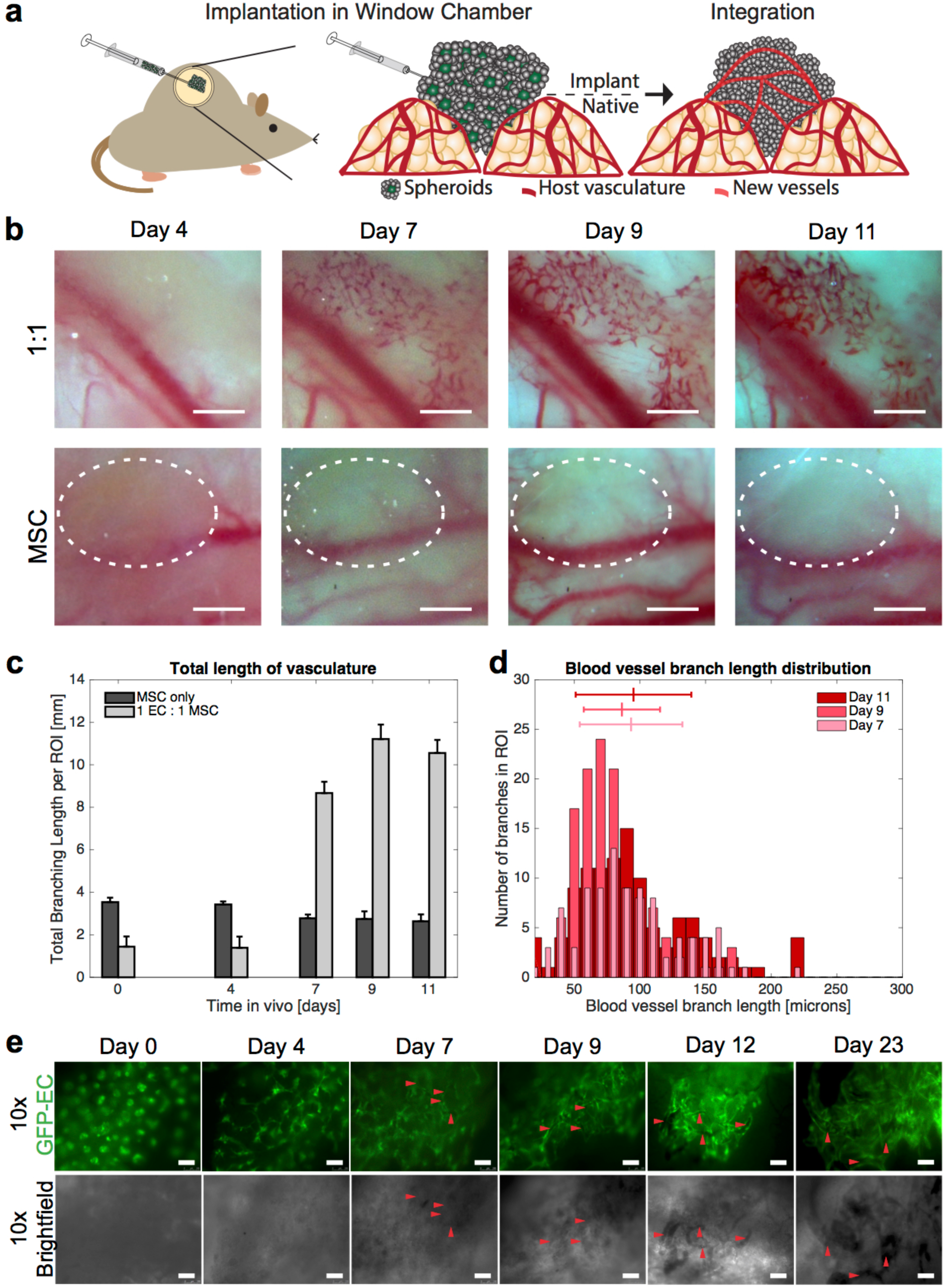
Rapid *in vivo* vascularization in healthy mice upon injection of organoids, as observed in real time via a window chamber. **(A)** Schematic diagram of experimental setup for observing vascular formation and integration with host vasculature *in vivo* in real time via a window chamber. Organoids (from human cells formed under dynamic culture conditions in 200 µm microwells yielding organoids 71±5 µm in diameter) were injected into a window chamber implant in a SCID mouse. **(B)** Real-time *in vivo* stereoscopic images of prevascularized microtisses with 1 GFP-HUVEC : 1 hAMSC (top row) and unvascularized organoids with hAMSC only (bottom row) through window chamber at different time points. Scale bars are 500 µm. In the top row, newly formed vessels are apparent within 4 days, and blood-filled vessels observed by day 7. In the bottom row, the dashed white line indicates the area of organoids implant and no neo-vascularization was observed. **(C)** Quantification of neo-vascularization of the prevascularized organoids as the total length of vasculature within three ROI. The total length of vasculature increases substantially after day 7 for prevascularized organoids. There is no substantial difference in total length of the vasculature for the unvascularized organoids. **(D)** Histograms of branching length in the newly formed microvasculature (b and c) at day 7, 9 and 11. Lines above histogram indicate the mean branch length and standard deviation for day 7, day 9, and day 11 as 93 ± 39 μm, 86 ± 29 μm, and 93 ± 44 μm respectively. **(E)** Real-time *in vivo* images of prevascularized organoids with endothelial cells in green. Red arrow heads point to luminous, blood-filled vessels (as indicated by dark lines in fluorescence images and dark areas of brightfield images). Scale bar is 250 µm.

We followed the formation of new vasculature by taking epiflourescent and stereoscopic images through the window chamber. Stereoscopic imaging (for example, of the 1 HUVEC : 1 hAMSC conditions) showed vessel formation between day 4 and 7 (Fig. 5b, top rows), with the implanted vasculature connected to the host vasculature and becoming perfused (Fig. 5b, top rows). After just 7 days, host perfusion of the implanted vasculature was prominent and intense. The vessels were functional for the remaining 16 days of the 23-day *in vivo* studies. Quantitatively, we measured the total length of perfused vasculature in three regions-of-interest (ROIs) within the area of injected organoids (Fig. 5c). At day 7, areas injected with pre-vascularized organoids showed significant formation of new perfused vasculature, while areas injected with organoids consisting of MSCs only showed no increase in perfused vasculature (Figs. 5b bottom row and 5c). For all four mice tested (each with multiple conditions in the window chamber), all conditions with EC-containing organoids showed rapid vascularization of the injected organoids.

We also explored whether this self-organizing, “micro-to-macro” strategy could provide a limited but reproducible level of architectural control in the overall branching length of implanted, perfused microvasculature. Specifically, we hypothesized that average distances between endothelial cores could be related to diameters of organoids. The mean length of the perfused branches for 1 EC:1 MSC at day 7 was 93±39 μm, with minimal changes by day 9 to 11, when the mean branch length was 86±29 μm and 93±44 μm, respectively (Fig. 5d). Indeed, the length of the newly formed vasculature’s branches reflected core-to-core distances between the densely packed, injected organoids with diameters of 71±5 μm.

We also used epiflourescence and confocal microscopy to characterize the formation and integration of the new vasculature. Observing the GFP-labeled HUVECs through the window chamber (Fig. 5e), we noticed the endothelial cores connecting with each other over time: the ECs initially appeared as discrete cores (day 0), then sprouted toward neighboring cores (day 4), connected with the host vasculature and became perfused (day 7), and stabilized as the perfused vascular network matured (day 9, 12, and 23). Between days 4 and 7, the network matured to form lumens (Fig. 5e, red arrows). (We further confirmed the lumenous structure of the newly-formed, perfused network on day 11 via confocal microscopy on day 11, Supplementary Video 4.) Moreover, we observed that areas indicating newly formed lumens (consisting of GFP-labelled HUVECs) co-localized with areas indicating host blood perfusion, further confirming that it was the newly-formed lumenous vasculature that was perfused, rather than angiogenesis from the host into the implanted tissue.

## Discussion

### Using sacrificial hydrogels to produce organoids with high reproducibility and scalability

Like the development of micromachining techniques for producing MEMS structures reproducibly and on a large scale, we have developed a technique to use sacrificial hydrogels to produce clusters of self-organized cell-based structures with high reproducibility and scalability. Previously, we and other groups have shown the use of microfabricated hydrogels, including sacrificial techniques, to form *in vitro* microvascular networks ^31,37,41,42^. This paper demonstrated the dynamic structure of hydrogels can also be exploited to produce and gently release organoids for cell therapy.

For purposes of cell therapy, it is critical for clinical efficacy, process control, and regulatory approval that cells introduced into the body are generated via tightly controlled processes and exhibit reproducible origin, size, and structure. Previous studies have observed that a “lack of control over the process is likely to underpin the variability in systems and experiments that, with few exceptions, does not allow [organoids] to yield their full potential”, and the importance of achieving reproducible “organoid size, shape, cellular composition and 3D architecture” in future research on organoids as well as use for therapeutic purposes ^12^. Compared to current organoid systems, our method can generate self-organized multicellular aggregates with both high yield (Supplementary Table 1) and high reproducibility over aggregate size and cellular organization (Supplementary Table 2). Moreover, the aggregate size and features of cellular organization can be tuned (Supplementary Table 2), as our method bears similarities to MEMS fabrication technologies (in contrast to “cells in gels” systems which feature a distribution of pore sizes). In this study, sizes and internal architectures of the organoids were reproducible for different types of cells (MSCs and ECs of mouse and human origin), cell ratios, and overall size of microwells which determined the diameter of the contracted organoids. Even at the tissue level both *in vitro* and *in vivo*, branching lengths of the vascular network were reproducible (by contrast, microtissues with ECs had previously yielded non-uniform branching lengths^7,16,43^.)

Also, an ideal method for generating organoids should be scalable and gentle. In the common hanging-drop method, 384 organoids could be produced in the area of an overall standard well plate (with the overall scalability limited by the number of wells ^14^), whereas the smallest construct shown in Fig. 2 produces 24,000 organoids in the same area with fewer steps needed (e.g. media-changing steps, one alginate dissolving step), all of which could be automated by liquid handling. The release of organoids is gentle even at a large scale, in contrast to vigorous pipetting or high-speed centrifugation for current microwell procedures. For cell therapy, it is important that the integrity of the cells be preserved (for example, a FDA guidance document points to the need “to preserve integrity and function so that the products will work as they are intended” ^44^. Beyond cell therapy, large-scale and effective production of organoids (beyond the quarter billion cells demonstrated) could also support studies in developmental biology, cancer cell intravasation ^16^, and organ printing.

### An advanced, controlled form of cell therapy

In past studies, needle injection (and organ printing) with unilaminar vascular organoids ^45,46^ had been challenging due to shear stress formation. It would be advantageous in cell therapy to be able to deliver the cells via minimally invasive injection rather than invasive surgery. Our method produced organoids which held up intact to shear stress during injections, even with high (25 to 30)-gauge needles (Supplementary Figure 9). This behavior may partially have been due to a shell of MSCs which protected the endothelial structure; interestingly, previous studies have also shown that the MSCs could act as an immune-suppressive shield for cell therapy in addition to providing angiogenic signaling ^47,48^.

A potential application of these organoids is to treat peripheral artery disease, the most severe form of which is critical limb ischemia (CLI) which can lead to amputations ^49^. To date, over 50 cell-therapy trials are at clinical stages for treating CLI. Many trials involve injecting MSCs ^49–53^ or ECs (such as MarrowStim)^52,54,55^, but the cells could die from deprivation of oxygen and nutrients before they are able to assemble into vascular networks *in vivo* and anastomose with host vasculature. The use of therapeutic pre-vascularized organoids could overcome many of the issues associated with currently cell therapy trials in clinical trials to treat CLI. In a clinical scenario, such an approach could be especially attractive for “no-option” patients on the verge of amputation with subsequently poor mortality outcomes (60% within five years of surgery ^49^).

## MATERIALS AND METHODS

### Experimental design

The objective of this study was to develop an approach to form self-organized organoids in a scaleable and gentle manner, for use as an injectable cell therapy. As such, we designed dissolvable alginate microwells to culture organoids, promote self-organization of the cells, and gently harvest the organoids. We then demonstrated their functionality in a healthy mouse model using a window chamber assay. All cells used in these studies were purchased commercially, all animal procedures were approved by the Columbia University Institutional Animal Care and Use Committee (IACUC) and all experiments were performed in accordance with relevant guidelines/regulations.

### Fabrication of alginate microwells

We developed an experimental setup to culture cellular organoids with high throughput and without the labor-intensive hanging drop approach. We seed the cells onto an alginate construct with between 379 and 30,000 microwells. The cells will settle into these microwells, and as the alginate provides no adherence structure for the cells, the cells will adhere strongly to each other forming spherical cell aggregates over the initial 24 hours.

The alginate microwells are cast on hydrophilic PDMS molds. We fabricated master molds in SU-8 (SU-8 3050, Microchem, Newton, MA) on 3-inch Si wafers (Silicon Sense, Nashua, NH) by photolithography as described before ^38^ to cast polydimethylsiloxane (PDMS, Sylgard 184, Essex Brownell, Fort Wayne, IN) replicas from the masters. We then made the PDMS molds hydrophilic by plasma treatment, and submerged them in distilled water to retain their hydrophilicity. We then autoclaved the PDMS molds prior to casting alginate.

We then prepared and autoclaved a 2% w/v alginate (Pronova UltraPure MVG, NovaMatrix, Drammen, Norway) or 7.5% w/v alginate (Alginic acid sodium salt, Millipore Sigma, St Louis, MO) in HEPES saline buffer solution (Ultrasaline A, Lonza, Basel, Switzerland). The alginate was pipetted into the PDMS molds. We used positive-displacement pipettes for accurate pipetting of viscous alginate solutions and to avoid bubbles. We closed the top of the molds with cellulose dialysis membranes (6000 Da MWCO), and flattened the membranes using the edge of a sterile glass slide. A 60 mM CaCl_2_ HEPES buffer solution was pipetted on top of the membrane for at least 60 min to crosslink the alginate at room temperature. We removed the hydrogels from the molds using sterilized tools, and placed the hydrogels in HEPES saline buffer solution (Ultrasaline A, Lonza, Basel, Switzerland) supplemented with 1.8 mM CaCl_2_ (to prevent leaching of the calcium ions from the hydrogels). We then transferred the alginate hydrogels into sterile culture ware, such as 24-well plates (Fisher Scientific, Fair Lawn, NJ) with the open micro wells facing up and stored them at 4°C until further use.

### Uncrosslinking of sacrificial alginate

To determine the length of time required to uncrosslink the microwells, 7.5% w/v alginate microwells were fabricated as described in the previous section, and stored in 1.8mM CaCl_2_ overnight. The following day, the alginate microwell scaffolds were transferred to preweighed, individually cut wells (from a 24 well plate), any excess CaCl_2_ was removed, and the initial mass of the alginate scaffolds was measured. We then added 1 mL of PBS, 0.5% w/v sodium citrate, or 5% w/v sodium citrate to the well, and after 1 minute the excess supernatant was removed, the remaining alginate scaffold was weighed, and a fresh solution of PBS, 0.5% sodium citrate or 5% sodium citrate was added. This was repeated until the alginate microwell scaffold was fully uncrosslinked (by the sodium citrate).

### Cell sources

GFP-labeled human umbilical vein endothelial cells (GFP-hUVECs) (Angioproteomie, MA, USA) were cultured in Endothelial Growth Medium 2 (PromoCell, Heidelberg, Germany). Adipose derived human mesenchymal stem cells (hAMSCs) (Promocell, Heidelberg, Germany) were cultured in Mesenchymal Stem Cell Growth Medium (Promocell, Heidelberg, Germany). RFP-labeled mouse mesenchymal stem cells (RFP-mMSCs) (Cyagen, CA, USA) were cultured in DMEM with 10% FBS and 1% PenStrep (all from LifeTechnologies). All cells were gently passaged at 80-90% confluency using TrypLE (LifeTechnologies) and used only until passage P6 and mMSC until P8. Cells were cultured in 37°C and 5% CO_2_-balanced, humidified atmosphere.

### Fabrication of organoids

HUVECs and MSCs were harvested from 2D cell culture, counted and desired cell ratios of HUVECs to MSCs were prepared: MSC only, 1 HUVEC to 3 MSC, 1 HUVEC to 1 MSC and 3 HUVEC to 1 MSC.

Then the 1.8 mM CaCl_2_ solution that the alginate microwells were stored in was removed, and replaced with DMEM (ATCC, Manassas, VA). The microwells were then placed in the incubator at 37°C and 5% CO_2_ to equilibrate for at least 20 minutes. Then DMEM was removed and the microwell constructs gently dried using surgical spears (Braintree Scientific, Braintree, MA) leaving the microwell features covered.

Cell suspensions were then pipetted on to alginate molds of 100, 200 and 400 µm microwell size using a positive displacement pipette. Cells were left to settle to the bottom of the microwells for 20 minutes and the culture wells were then filled up with culture medium.

### Culture media

The first 3-4 days after seeding, the cells were cultured in maintenance medium: Dulbecco’s Modified Eagle Medium (DMEM) with 10% Fetal Bovine Serum (FBS) and 1% PenStrep (all from LifeTechnologies), with 50 μg/mL Sodium L-ascorbate (Sigma-Aldrich). For Fig. 4 and 5, the maintenance medium also included 20 mM Hepes (Fisher Scientific, Fair Lawn, NJ), 1 μM Insulin (LifeTechnologies, Carlsbad, CA), 250 nM T3, 1 μM dexamethasone, 0.5 mM IBMX, 50 μM Indomethacine, 1 μM Rosiglitazone and 1 μM CL316243 (all from Sigma, St. Louis, MO). After the first 3-4 days, the media was changed from maintenance medium to vasculogenic medium: Dulbecco’s Modified Eagle Medium (DMEM) with 10% Fetal Bovine Serum (FBS) and 1% PenStrep (all from LifeTechnologies), with 50 μg/mL Sodium L-ascorbate (Sigma-Aldrich), 40 ng/mL bFGF and 40 ng/mL VEGF recombinant human protein (both from LifeTechnologies). For Fig. 4 and 5, the vasculogenic media also included 20 mM Hepes, 1 μM Insulin, and 250 nM T3. The cells were cultured in vasculogenic medium up to day 11. Cell media was changed every other day.

### Harvesting of organoids

To collect organoids, the alginate hydrogel was uncrosslinked ^37^. For this, the culture medium of the organoids was replaced with 5%w/v sodium citrate solution for approximately 20 min. This chelator liquefied the alginate, and allowed for resuspension of the organoids in a desired medium. Organoids were then centrifuged at 300 rpm for 5 minutes or as specified and the organoid pellet carefully collected for further use.

### Organoid fusion

To test the ability of organoids to assemble *in vitro* and fuse to a larger tissue, 200 μm sized organoids of only hAMSCs, 1 EC : 3 hAMSC and 1 EC : 1 hAMSC ratio were placed in a 400 μm sized microwell of collagen-doped alginate ^38^ composed of 3.5% collagen and 1% alginate. These organoids had previously been prevascularized as described above. Organoid fusion was conducted in vasculogenic medium and observed for 24 hours.

### Formation of macrotissue

To yield a large enough number of organoids in parallel to produce a macroscopic tissue, we seeded a co-culture of 1 MSC : 1 EC ratio onto an alginate microwell construct that fits into a 60 mm culture dish and produces over 30,000 organoids (Fig. 2a and c). The cells were cultured in maintenance medium without growth factors for 4 days with daily media change due to the high number of cells. The organoids were collected by removing the medium and uncrosslinking the alginate with 5 mL 5%w/v sodium citrate solution. The alginate liquefied and the organoid solution was gently collected, spun down and resuspended in 1 mL vasculogenic media. To facilitate sustained culture and imaging of the macro-tissue, we had constructed a 1 cm^2^ cylindrical hole in a 1 cm thick 10% agarose layer in the middle of a 60 mm dish. The hole was made by placing a 1 cm^2^ by 1 cm PDMS mold in the middle, pouring on agarose and removing the PDMS cylinder when the agarose had gelled. The 1 mL organoid suspension was pipetted into the hole, allowed to settle for 1 hour, and then had 5 mL vasculogenic media added on top. The media was changed daily.

### Window chamber surgery

Organoids were collected as described above. The suspension was gently spun down at 220 g for 5 minutes. The supernatant was removed and the organoids resuspended in 200 µl PBS. In vitro created, prevascularized and non-prevascularized organoids were implanted in a window chamber to allow for continuous in vivo monitoring of the vascularization and integration process. Window chamber surgeries were conducted as described previously ^56,57^. A titanium window chamber (APJ Trading, Ventura, CA) was surgically implanted midline on the dorsum of male SCID mice (strain: ICRSC-M-M, 5-6 weeks of age, Taconic, Hudson, NY) after hair removal and ethanol and iodine disinfection.

To reduce variability between mice, prevascularized and non-prevascularized organoids were implanted in individual compartments of the same window chamber. Organoids were delivered by injection and pipetting underneath the fascia of connective tissue to the subcutaneous adipose tissue. Window chambers were closed with a circular glass cover slip and a retaining ring (APJ Trading, Ventura, CA). A custom-made 3D printed window chamber backing was attached to reduce skin movement in the window chamber. In a subset of experiments, a custom-made ultem plastic 9 well array was screwed onto the front frame of the window chamber to allow for high throughput in vivo testing. Here, organoids were placed into one of the 9 wells.

Animals were housed aseptically in frog cages to allow for enough clearance for the window chamber while still permitting easy access to standard laboratory chow (Irradiated globle rodent diet, Fisher Scientific, Fair Lawn, NJ) and drinking water ad libitum. Follow up buprenorphine administration (0.1mg/kg bodyweight) for pain management was given subcutaneously every 6-12 hours after surgery for the next 2 days post-OP. CO_2_ euthanasia and cervical dislocation were performed after 30 days or earlier if necessary.

The animal procedures were approved and carried out in accordance to local regulations and authorities. The surgeries were conducted in aseptic technique.

### Imaging

A Leica DMI 6000B inverted microscope with 4x and 10x objectives, equipped with a motorized stage (Leica Microsystems, Bannockburn, IL) and a QImaging Retiga 2000R monochrome camera (QImaging, Surrey BC, Canada) was used to acquire fluorescence and brightfield images. Leica LAS X software was used for image acquisition. Cropping, color adjustments and contrast enhancements of images as well as Z-stack projections were performed in ImageJ. For time lapse imaging of organoid formation and organoid fusion an environmental chamber was used to maintain 37°C and 5% CO_2_. Images were acquired every 30min. Confocal images of the window chamber were taken using a Leica SP5 confocal system with a 10.0x 0.30 N.A. objective. To be able to image the window chamber mice were anesthetized with isoflurane. Due to the stressfulness of the anesthesia of the imaging procedure, in vivo images were acquired every 2-3 days.

To precisely observe individual organoids, we took stacks of confocal images (1024×1024 pixels, 41 images with a z-spacing of 0.25 microns) using a Leica SP5 confocal microscope, with a 100x 1.43 N.A. oil immersion objective (Leica Microsystems) at a resolution of 0.132 µm/pixel (image stacks were thus 135 mm * 135 mm * 10 mm). We simultaneously collected the differential interference contrast (DIC) images as well as the RFP- and GFP-signal.

## Acknowledgements

We acknowledge technical assistance by Yaas Bigdeli and Ayse Karakecili, and Mohammed Shaik and Elizabeth Hillman for help with imaging.

## Author contributions

N.S.R., R.z.N., B.M.G., and S.K.S. conceived the project and designed the experiments. N.S.R., P.N.A., R.z.N., K.L., W.L., C.P., C.H., Q.F., Z.S., R.P.S.-M., J.C., J.E.G., N.S., T.H. and B.M.G. conducted the experiments and analyses. N.S.R., P.N.A., R.z.N. and B.M.G. analyzed and interpreted the data. N.S.R., P.N.A. and R.z.N. prepared the figures, and N.S.R., R.z.N., and S.K.S. wrote the manuscript with contributions from P.N.A. and B.M.G.. N.S.R., B.M.G., P.N.A. and S.K.S. supervised the project. All authors have reviewed the manuscript.

