## Supplementary Materials for "Injectable therapeutic organoids using sacrificial hydrogels"

**SUPPLEMENTARY FIGURES**

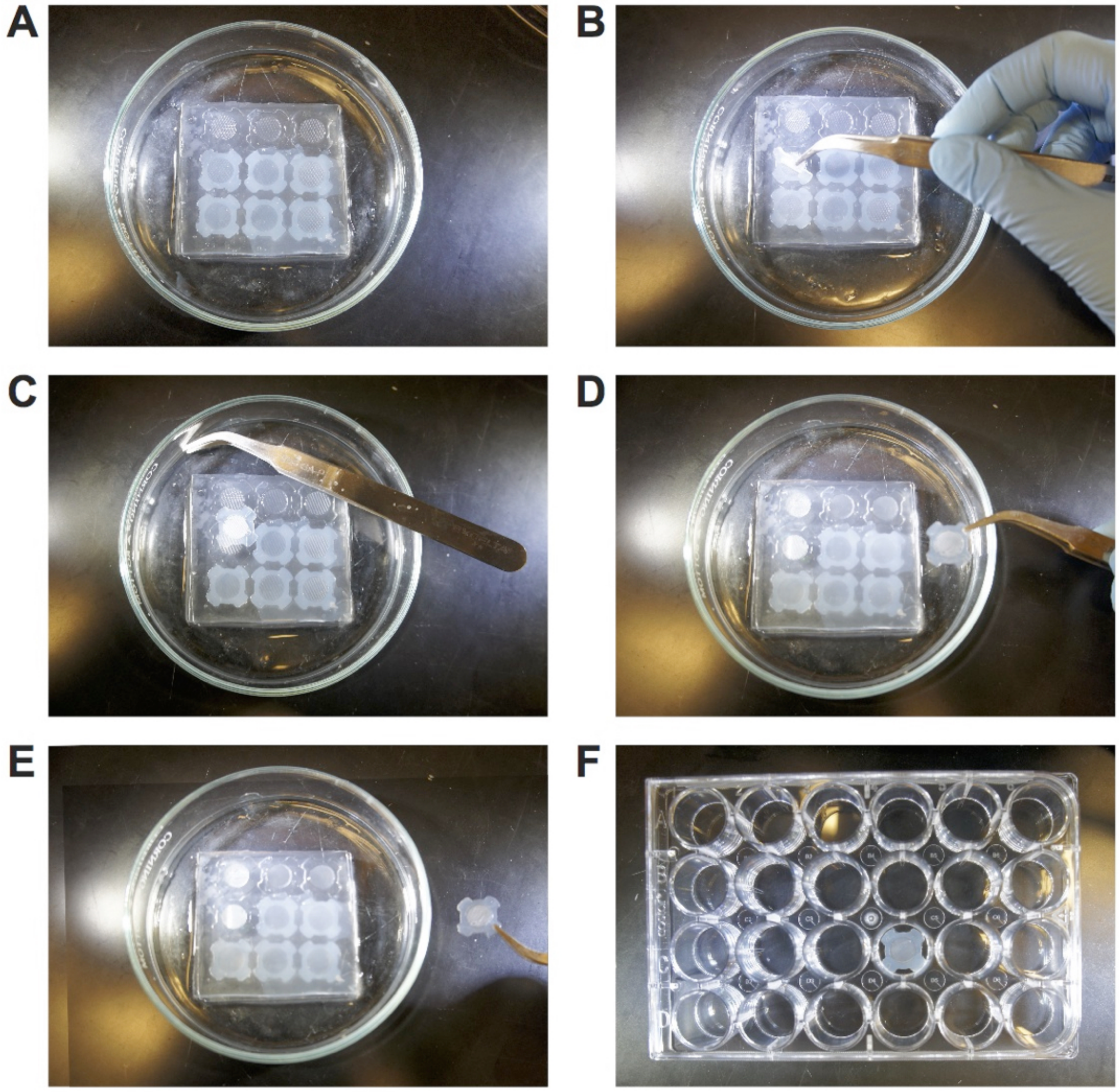


**Supplementary Figure 1. Fabrication of alginate-based micro-wells. (A)** shows six cross-linked alginate microwell constructs in the PDMS mold with the top three constructs removed. **(B-D)** The alginate molds are carefully loosened with a pair of sterile tweezers and removed from the mold holding on only to the “wings” to preserve the microwell structures in the middle. **(E)** The alginate constructs must be flipped to expose the microwells before being placed in the well plate **(F)**.

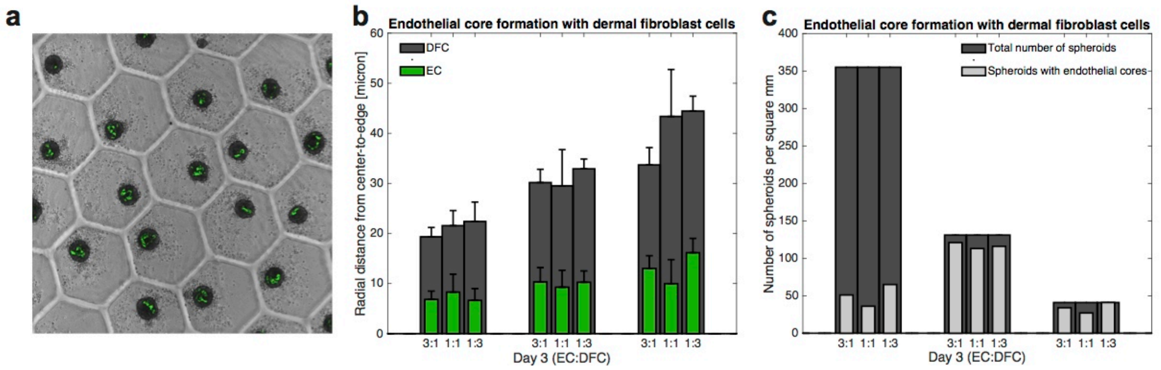


**Supplementary Figure 2. Endothelial core formation in co-culture with HUVECs and human dermal fibroblast cells (hDFCs). (A)** Image of organoid and endothelial core formation of 1 EC : 1 DFC in 400 μm wells after three days culture in maintenance medium. **(B)** Barplot of the radius of the smallest circle that can contain all ECs (green) or all DFCs (dark gray) for the different well sizes at Day 3. **(C)** Barplot of the number of organoids/spheroids formed per mm$^2$ (dark grey) and the number of those organoids that contain endothelial cores (light grey).

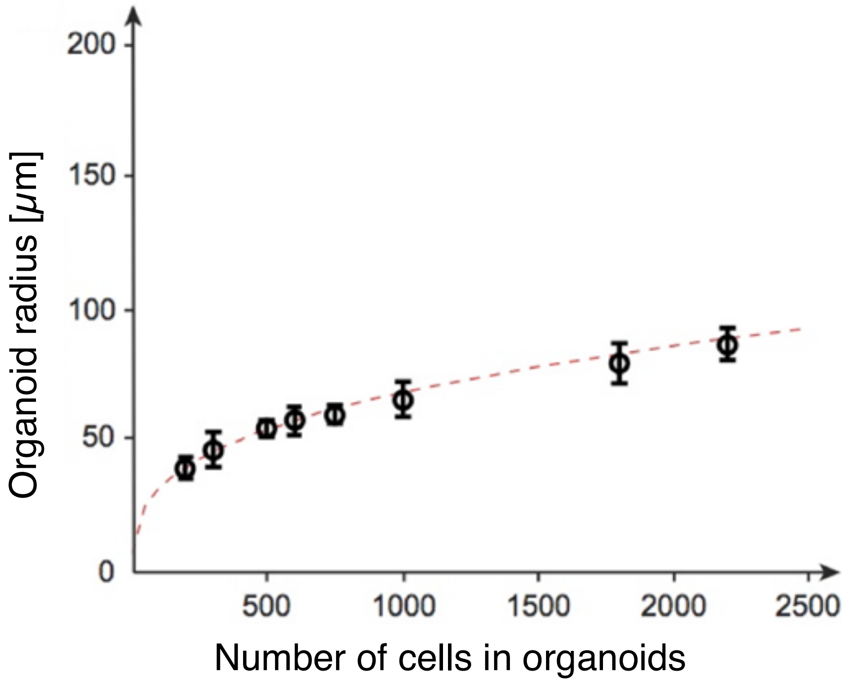


**Supplementary Figure 3. Organoid size is a function of the number of cells inside.** Our method provides two mechanisms to control the number of cells in each organoid; either by changing the size of alginate wells or by changing the seeding cell density. Utilizing both, we got organoids with 200, 300, 500, 600, 750, 1000, 1800, and 2200 cells. The radius of the resulting, fully-aggregated organoids after 3 days follows the expected equation $r_{organoid} = (6/\pi\ v_{\mathrm{cell}}\ n_{\mathrm{cell}})^{1/3}/2$.

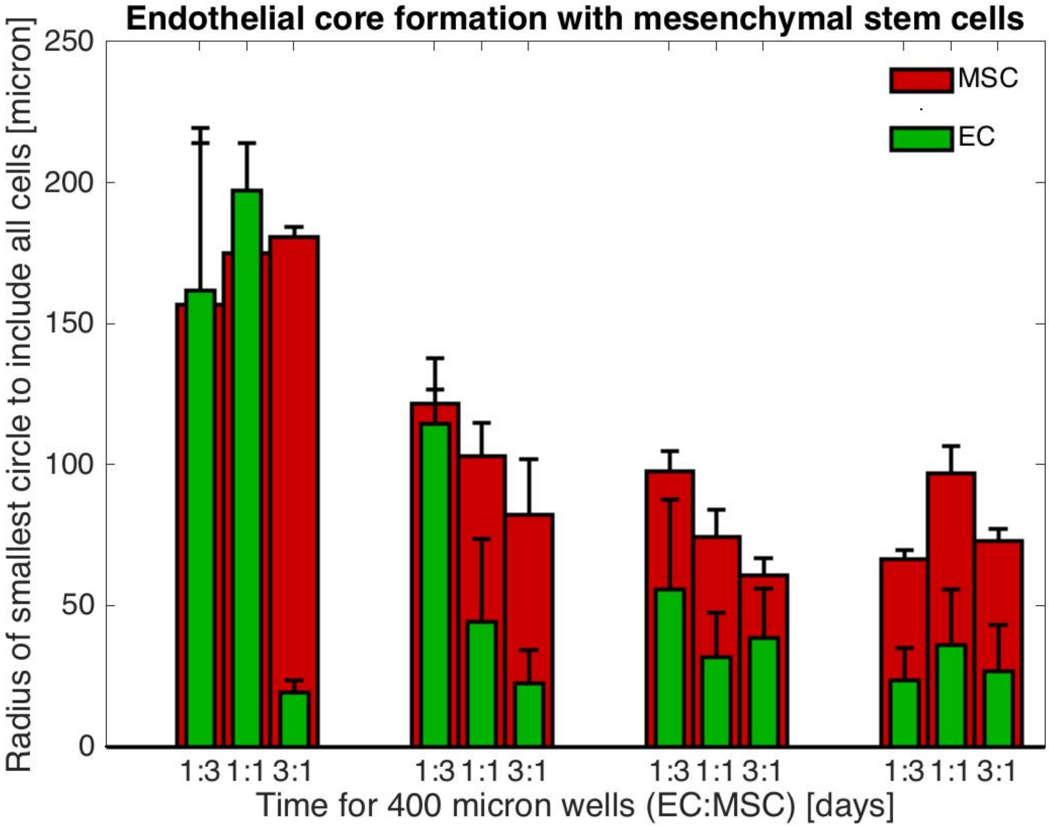


**Supplementary Figure 4. Organoid contraction and endothelial core formation in 400-micron microwells from day 0 to day 3.**

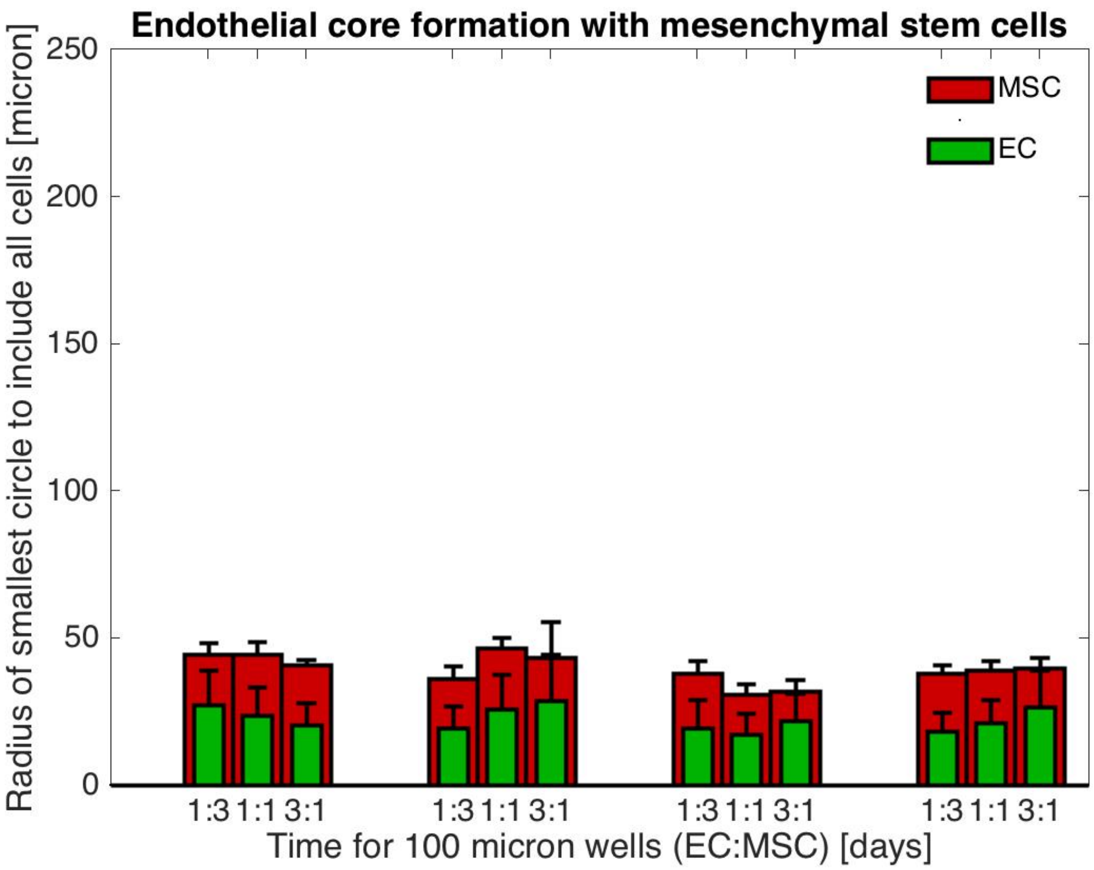


**Supplementary Figure 5. Organoid contraction and endothelial core formation in 100-micron microwells from day 0 to day 3.**

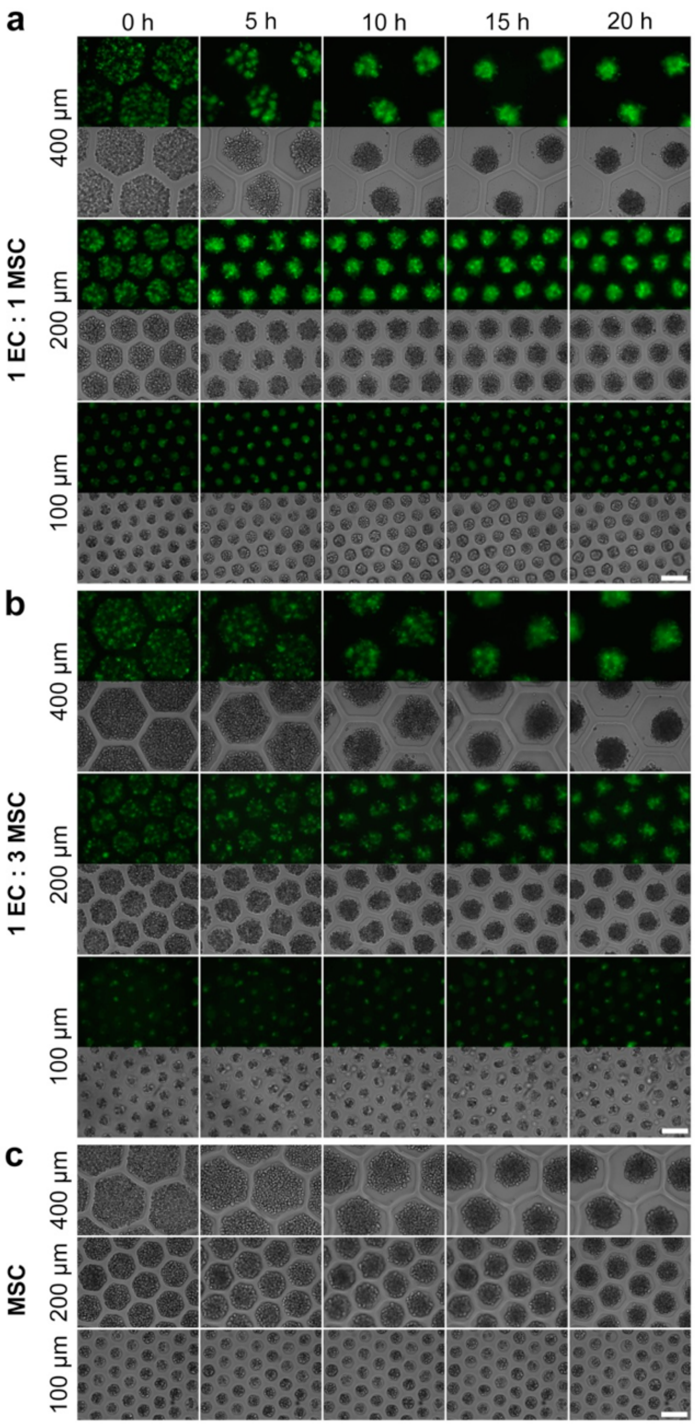


**Supplementary Figure 6. Initial phase of organoid formation and self-sorting.** Shown are the first 20 hours of self-assembly and self-sorting immediately after seeding GFP-HUVEC and hAMSC cell mixtures in alginate microwells (100, 200, 400 µm). Fluorescent and brightfield images were acquired with a 10x objective. Scale bars are 200 µm. **(A)** 1 HUVEC : 1 hAMSC organoids. **(B)** 1 HUVEC : 3 hAMSC organoids. **(C)** hAMSC only organoids.

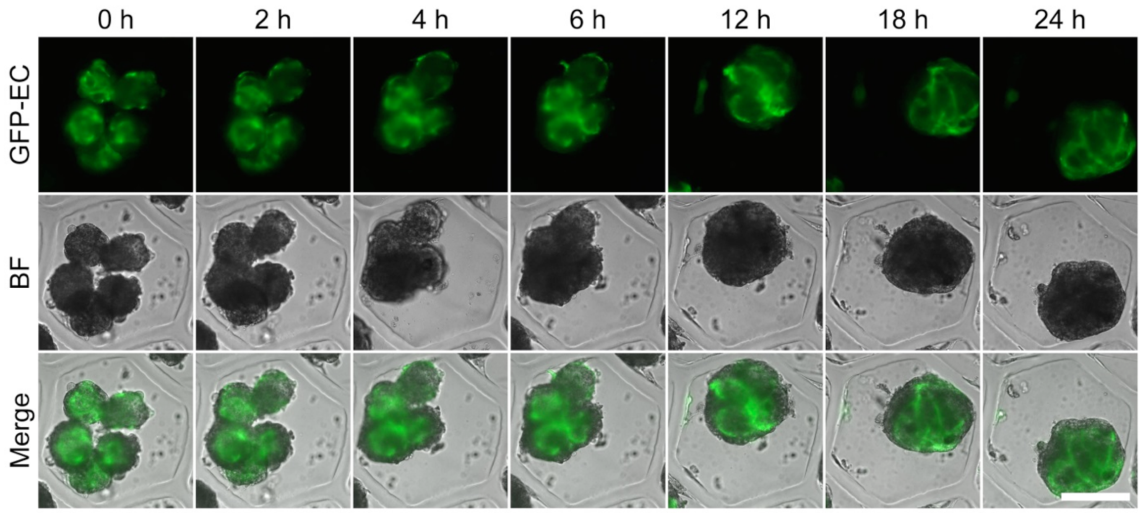


**Supplementary Figure 7. Organoid fusion to form macrotissue.** Organoids were created in 200 μm sized microwells at 1:1 and 1:3 GFP-HUVEC to hAMSC ratio. Organoids were cultured for 3 days in maintenance medium and 6 days in a vasculogenic medium to induce prevascularization (as seen at 0 h). For this assay the same vasculogenic medium was used. Prevascularized organoids were placed in a collagen-alginate microwell. GFP fluorescence and brightfield images were acquired with a 10x objective in time-lapse microscopy over 24 h. Scale bar is 200 μm.

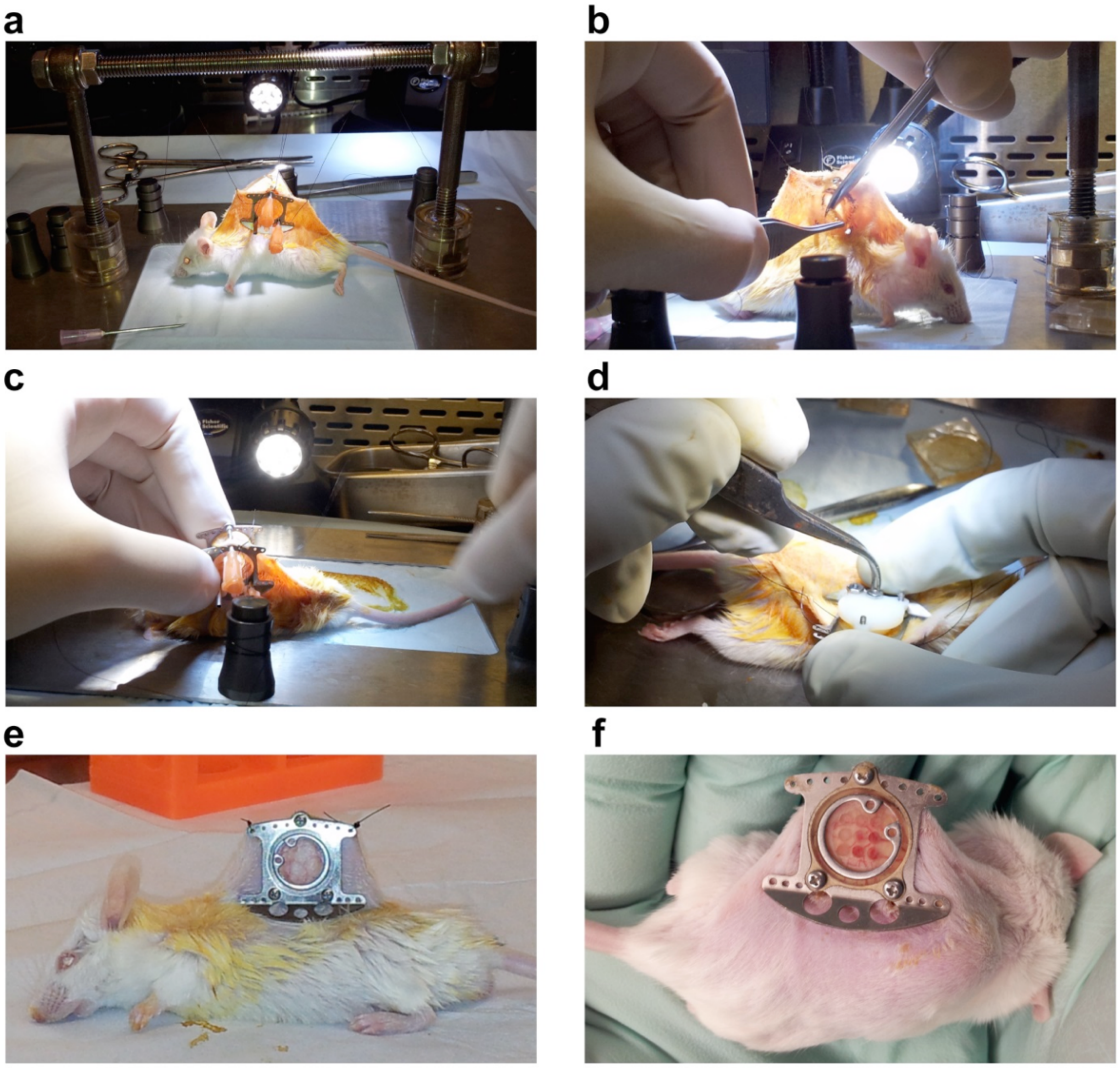


**Supplementary Figure 8. Window chamber surgery. (A)** Dorsal skin was spread out and backside of window chamber was connected. Needles were used to create holes for window chamber. **(B)** A circle was marked on the skin to indicate the opening for the window chamber and the skin was carefully cut out. **(C)** Front and back of window chamber were assembled. **(D)** Nuts were used to lock the window chamber. A custom 3D printed backing was used to hold the skin in place. **(E)** Closed window chamber in place; Secured with lateral sutures. **(F)** A custom-made window chamber insert to test up to 9 conditions simultaneously was included.

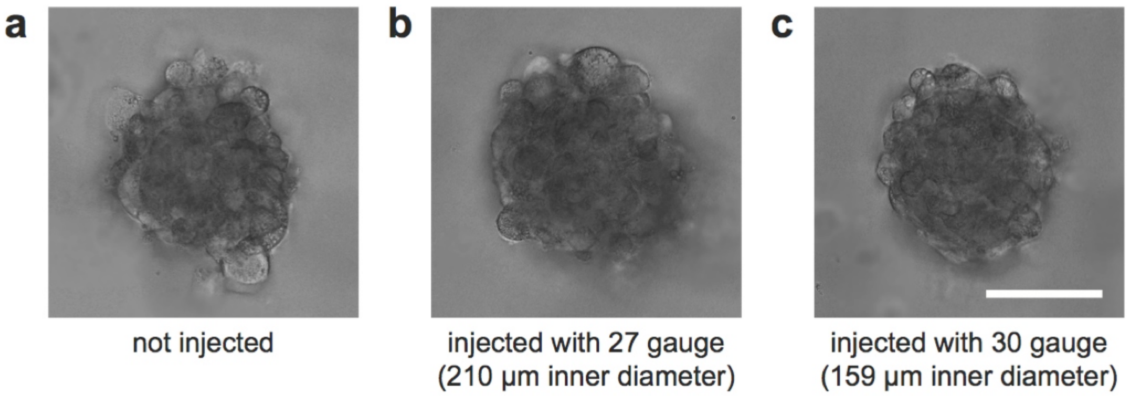


**Supplementary Figure 9: Robustness to shear stress.** Images of organoids before **(A)** and after injection through a 27 gauge needle **(B)** and a 30 gauge needle **(C)**. Scale bar is 50 µm.

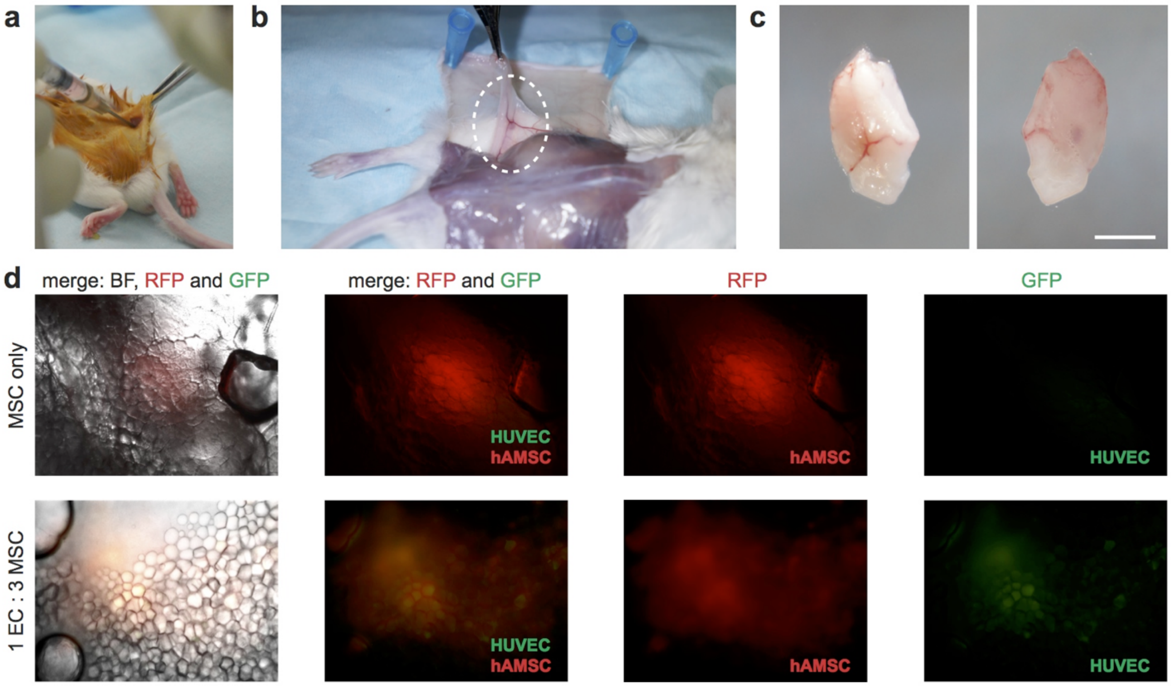


**Supplementary Figure 10. Procedure for injecting organoids *in vivo* (without window chamber).** (**A**) Injecting a solution with organoids (1EC:3MSC and 200 µm size) into the adipose tissue of the obliques. A small incision along the back was made to ensure the correct position of the injection site. (**B**) The mice were sacrificed and the entire obliques (white dashed circle) were excised after 18 days of implantation. (**C**) Front and back of the extracted obliques. Scale bar is 2.5 mm. (**D**) Merged and individual images of brightfield and fluorescent images showing the position of the injected organoids. Top: RFP-MSC only organoids. Bottom: 1 GFP-ECs : 3 RFP-MSC ratio organoids.

**Supplementary Table 1. Table of advantages for different cell aggregation production methods**

| Cell Aggregate Production Method | High throughput | Output (no. of cell aggregates) | Easy automation (liquid handling in all steps) | Size control | Continuous observation | Easy harvesting | Biologically inert process suitable for *in vivo* use | Reference |
|---|---|---|---|---|---|---|---|---|
| Sacrificial hydrogel microwells | ++ | 10,000s per mold | + | + | + | + | + | Current Study |
| Hanging drops | - | 60 to 384 per culture plate | + | + | - | + | + | [14,58,59] |
| Spinner culture | + | 1,000s per spinner flask | + | - | - | + | + | [13] |
| Microwells | + | 1,000s per mold | - | + | + | - | +<br>- (for PDMS microwells) | [1,9,35,60,61] |
| Non-adhesive 96 well plate | - | 96 per culture plate | - | + | + | - | + | [16,18,19] |

**Supplementary Table 2. Reproducibility and control of vascularized organoids**

| Reference | Method for producing vascularized organoids | Control over size in cited study | Structural organization of cells within organoid | | Control over shape (circularity) | Organized vascular architecture within aggregates |
|---|---|---|---|---|---|---|
| | | | Observation of core-shell formation | Control over core-shell structure | | |
| Current study | Sacrificial hydrogel microwells | Yes<br>-by controlling size of alginate microwells | Yes | Yes<br>-by changing presence of growth factors<br>-by controlling ratio of ECs to MSCs | High | Present |
| Kelm et al.[59] | Hanging drop | Yes<br>-by controlling cell number | Yes | Not demonstrated | High | Present |
| De Moor et al.[1] | Microwells | No | Yes | Not demonstrated | High | Present |
| Wenger et al.[19] | Non-adhesive 96 well plate | No | Yes | Not demonstrated | Medium | Absent |
| Kim et al.[62] | Thermoresponsive hydrogel sheets | Yes<br>-by controlling size of hydrogel sheet | Yes | Yes<br>-by controlling method of seeding cells on hydrogel surface<br>-by controlling HUVEC cell density | Medium | Absent |
| Tiruvanna-malai et al.[63] | Cells embedded within gels | No | No | No | Low | Present |

### MOVIES

**Supplementary Movie 1. Dissolution of alginate as sacrificial scaffold for gentle release and harvesting of organoids.**

**Supplementary Movie 2. 3D formation (self-assembly and self-sorting) of organoids over the first 20 hours.** Subsets of data are shown in Fig. 3b and Fig. S6a.

**Supplementary Movie 3. Fusion of organoids of different EC:MSC ratios. Shown are organoid ratios 1:1, 1:3 and MSC only.** Subsets of data are shown in Fig. 3c and Fig. S7.

**Supplementary Movie 4. Spatially controlled organoids *in vivo* at day 11.** Z-stack video of organoids in window chamber with 9-well insert at day 11 (maximum intensity projection is shown in Fig. 1a).
